# Reconstructing and characterizing focal amplifications in cancer using AmpliconArchitect

**DOI:** 10.1101/457333

**Authors:** Viraj Deshpande, Jens Luebeck, Mehrdad Bakhtiari, Nam-Phuong D Nguyen, Kristen M Turner, Richard Schwab, Hannah Carter, Paul S Mischel, Vineet Bafna

**Affiliations:** Department of Computer Science and Engineering, University of California at San Diego, La Jolla, California 92093, USA, United States; Bioinformatics and Systems Biology Program, University of California at San Diego, La Jolla, California 92093, USA, United States; Ludwig Institute for Cancer Research, University of California at San Diego, La Jolla, California 92093, USA, United States; Department of Medicine, Division of Hematology-Oncology, School of Medicine, University of California at San Diego, La Jolla, California 92093, USA, United States; Department of Medicine, Division of Medical Genetics and Moores Cancer Center; University of California at San Diego; La Jolla, CA 92093; USA, United States

## Abstract

Focal oncogene amplification and rearrangements drive tumor growth and evolution in multiple cancer types. We developed a tool, AmpliconArchitect (AA), which can robustly reconstruct the fine structure of focally amplified regions using whole genome sequencing. AA-reconstructed amplicons in pan-cancer data and in virus-driven cervical cancer samples revealed many novel insights about focal amplifications. Specifically, the findings lend support to extrachromosomally mediated mechanisms for copy number expansion, and oncoviral pathogenesis.

---

Cancer is marked by somatic DNA lesions. While these include small nucleotide changes, and chromosomal aneuploidies, focal amplifications of smaller regions are also a prominent signature in a large proportion of human cancers^1^. Focally amplified regions are found to be hotspots for genomic rearrangements, which can include the juxtaposition of segments of DNA from distinct chromosomal loci, into a single amplified region^2–8^. While common, these types of focal gene amplification present a mechanistic challenge - how do multiple regions from one or more chromosomes rearrange together in cancer? Our team recently showed that focal amplification in nearly half of samples across a variety of cancer types can be explained by circular, extrachromosomal DNA (ecDNA) formation^9^. Furthermore, ecDNA formation can dramatically change the outlook for tumor evolution even as compared other types of somatic mutations. Due to this renewed understanding, there is an urgent need for tools to study the biological properties of ecDNA, and more importantly facilitate ecDNA-based techniques for cancer treatment and diagnostics. Specifically, tools to elucidate the structure of ecDNA can provide insights into the mechanisms of oncogene amplification and evolution through complex rearrangements.

An ‘amplicon’ is defined as a set of non-overlapping genomic ‘intervals’ connected to each other and amplified, and an ‘amplicon structure’ as an ordered arrangement of the genomic ‘segments’ within the amplicon. An amplicon interval may be partitioned into multiple genomic segments depending on the rearrangement breakpoints within the amplicon structures. We recently found that oncogenes amplified on ecDNA are often part of highly rearranged amplicons, that may juxtapose regions from different chromosomes. Traditional structural variant (SV) analyses cannot decipher complex rearrangements^10–13^. The few methods that extend the analysis, chain together breakpoints into paths and cycles, but often do not reconstruct the amplicon in the specific region of interest, and do not provide a comprehensive view of alternative structures^14–19^. Reconstruction remains challenging due to extreme variability in copy counts (5X-200X) and sizes (100kbp-25Mbp) of amplicons, samples containing heterogeneous mixture of multiple amplicon structures, and inaccuracy of SV identification.

We describe AmpliconArchitect (AA), a tool for reconstruction of ecDNA amplicon structures using whole genome sequencing data (Fig 1A-J, Methods 1,2) that overcomes these difficulties by providing a versatile representation of an amplicon that encodes all supported structures and provides a framework for algorithmic reconstruction of possible structures.

**Figure 1.**
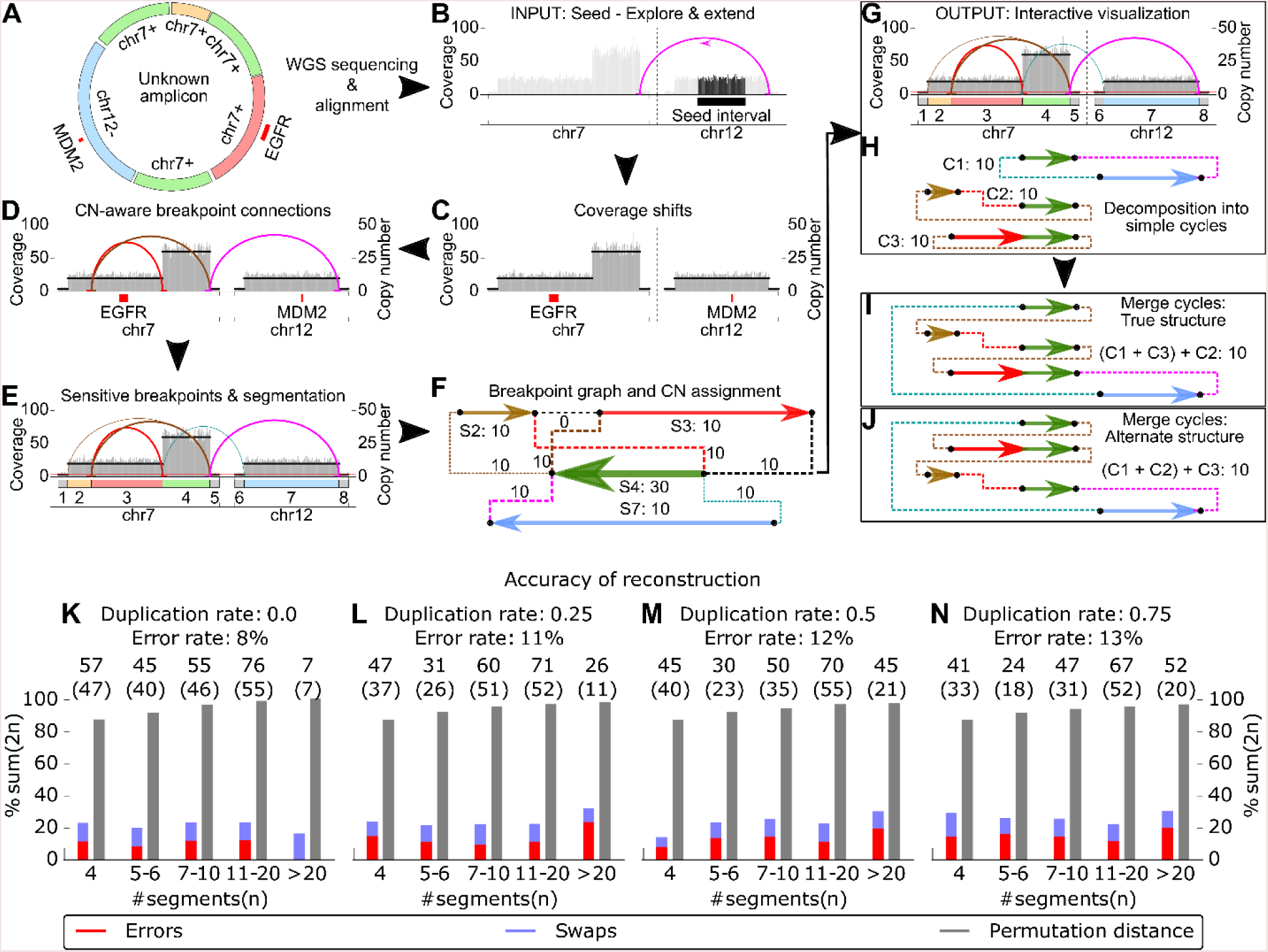
Schematic of Amplicon Architect (AA). AA takes as input: (A) aligned whole- genome sequencing data from a sample with an amplicon, and B) a ‘seed’ interval from the amplicon. It automatically searches for and identifies other intervals that are part of the same amplicon; C) Next, AA identifies breakpoints by segmenting intervals at positions with a sharp change in copy number, or (D) containing a cluster of discordant paired-end reads. Finally, (E) AA refines breakpoint locations. (F) The collection of segments and breakpoints is used to generate a breakpoint graph, and a balanced flow approach is used to refine segment copy numbers. (G, H) The entire graph describes a breakpoint signature and a succinct “SV view” of the amplicon, which is also decomposed into short basis cycles in the “Cycle view”. (I, J) Alternative merging of the short cycles with overlapping segments can generate multiple amplicon architectures consistent with the short-read data. (K-N): Amplicon reconstruction on a variety of simulations showed high fidelity of reconstruction (red bar, 11% error) relative to a random ‘permutation predictor’ (grey bar, 85% error). Swaps (blue bars) represent cases with alternative structures supported by the data. A high fraction of samples were reconstructed perfectly (parenthesized values), although performance decayed slightly upon increasing duplications in the amplicons. Values are presented as percentage of twice the sum of number of segments.

AA takes as input, short, paired-end reads mapped to the reference genome as well as a seed interval in an amplicon. It automatically searches for other intervals participating in the amplicon (Fig 1A-B), and then performs a carefully calibrated combination of copy number variant (CNV) analysis^20^ and SV analysis (Fig 1C-E). For algorithmic prediction of the amplicon structures, AA uses SV signatures (e.g. discordant paired end reads and CNV boundaries) to partition all intervals into segments and build a breakpoint graph. It assigns copy numbers to the segments by optimizing a balanced flow on the graph^21^ (Fig 1F). As short reads may not span long repeats, they cannot disambiguate between multiple alternative structures. AA addresses this in two ways. First, it creates a succinct “SV View” displaying the raw SV signatures including coverage depth, copy number segments and discordant genomic connections (Fig1G), which by itself is informative to the user for manual interpretation of the amplicon structure. Second, AA decomposes the graph into simple cycles, and provides a “Cycle View” to intuitively visualize the segments of the cycles in the context of the SV view, showing their genomic locations. The AA Cycle View provides a feature to interactively merge the simple cycles and explore candidate amplicon structures (Fig 1I-J, Supplementary Fig 1).

A robust amplicon reconstruction tool should predict the amplicon structure for a diverse set of focal amplifications observed in cancer in a high-throughput manner. Previous studies examining complex rearrangements tested small sets of cancer samples and validated individual rearrangements using PCR. On those samples, AA performed at least well as the other studies and identified SV signatures that other methods were unable to identify (Methods3, Supplementary materials, figshare). A complete validation of entire structures would require ultra-long read fragments (Mbp) or isolation of amplicons from the rest of the genome. Moreover, multiple experiments would be needed to test AA effectively on the diversity of reconstructed amplicon structures (e.g., see Fig S7, figshare). Therefore, we developed a simulation-based benchmarking strategy and error model to quantify the accuracy of predicted structures.

We simulated a diverse set of ~1000 amplicons, including rearrangements with varying levels of copy number (4x-32x), size (40kbp-2.4Mbp), number of rearrangements (0-16), duplication probability (0-0.75), and sequencing coverage (1x-32x) (Methods 4). AA had consistent performance in terms of prediction of SV signatures with changing copy number and coverage (Supplementary Fig 3). To measure the accuracy of the complete reconstruction, we developed a novel metric based on a ‘graph edit distance’, described informally by the number of operations required to transform the predicted amplicon structure into the true structure (Methods 5, Supplementary Fig 2). The metric partitions the graph operations into two categories: (i) number of *errors* caused by AA and (ii) number of swaps across *repeat branches* indicative of the number of cycle merging operations a user would need to perform to obtain the entire structure. The number of errors was normalized by the total segments in the amplicon and contrasted against a naive ‘permutation predictor’ which picks a random order of the amplicon segments from the copy number profile (Methods 6). This metric reported that even on our wide-ranging simulations, AA had an error rate <=11%, averaging about 1 error per 9 rearrangements (Fig 1K-N, Supplementary Fig 3, 4).

We applied AA to sequencing data of 117 cancer samples (sample set 1) (supplemented by 18 replicates and drug-treated variants) from 13 cancer types^9^ (Methods 7, Supplementary Table 1). We used the CNV tool ReadDepth^22^ (Methods1, Supplementary Table 2) to identify 255 focally amplified intervals in 55/117 samples with size >100kbp and copy number >5X. Using the 255 intervals as seeds, AA reconstructed 135 non-overlapping amplicons, each containing one or more seeds (Supplementary Table 3, figshare). We observed a range of amplicon structures including simple cycles, heterogeneous mixtures of related structures, a breakage-fusion-bridge amplification, and highly rearranged amplicons with intervals from one up to 3 chromosomes (Fig 2A-C, Supplementary Table 3, figshare). The number of detected rearrangements (breakpoint edges) per amplicon ranged from 0 to 49 with average of 4.9 rearrangements per amplicon, a likely underestimate, because of the low-coverage sequencing data.

**Figure 2.**
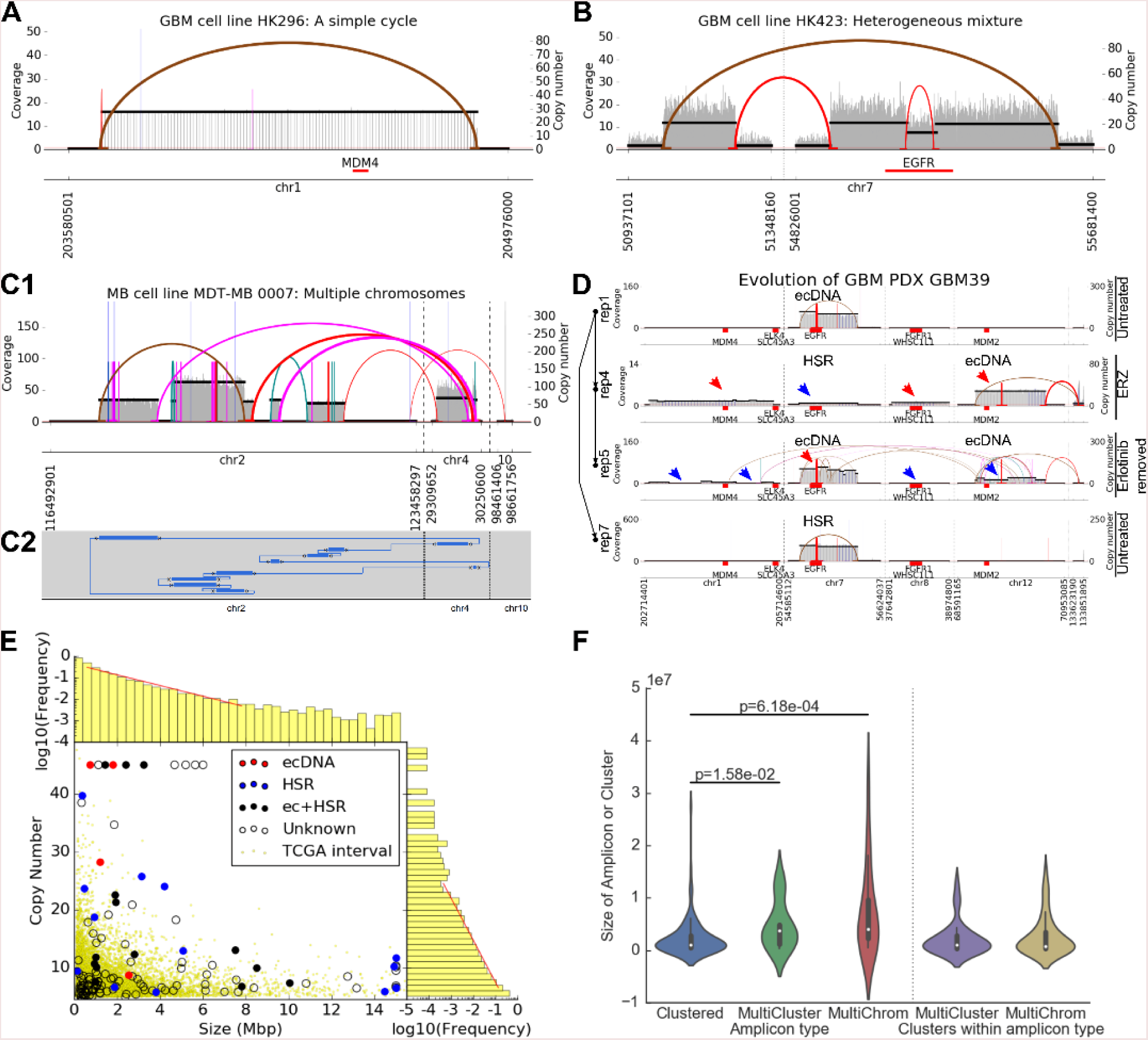
SV view of AA reconstructions. AA reconstructed 135 amplicon structures from 255 seed intervals in WGS of 117 cancer samples. The SV View of reconstructed amplicon structures shows (A) simple cycles; (B) heterogeneity with amplicons containing EGFR VIII deletion as well as the intact EGFR; and (C1) complex rearrangements such as a medulloblastoma multichromosomal amplification. (C2) A cycle view of the MB amplification. (D) A combined SV view of all amplicons in a GBM patient derived xenograft (PDX) evolving with time and in response to drug treatment shows interconversion of ecDNA and HSR, copy number changes (red arrows: increase, blue arrows: decrease) and structural changes. Left axis: passage of PDX, right axis: replicate state (ERZ: erlotinib resistant). (E) AA amplified intervals compared against 12162 amplified TCGA intervals shows significant overlap. The interval sizes (mean 1.74 Mb) and copy numbers (mean 3.16 copies) are exponentially distributed, with no clear distinction between HSR and ecDNA amplicons. (F) Amplicons with intervals from multiple genomic regions from the same chromosome (MultiCluster) and multiple regions from multiple chromosomes (MultiChrom) are larger in size that amplicons with all intervals in a single region on a single chromosome (Clustered), but size of amplified region within each cluster follows a similar distribution as the Clustered amplicons.

Typical mechanisms invoked for CN amplification rely on repeated breaks at fragile sites followed by duplication events. We used AA to test an alternative model, that (a) starts with breaks at random sites, followed by ecDNA formation; Poisson random breaks result in an exponential distribution of segment lengths. (b) Aggregation of ecDNA may create larger, highly-rearranged, multi-interval, structures. (c) Replication and independent segregation of ecDNA create cells with a diversity of copy numbers; however, (d) positive selection for higher copy numbers due to proliferative elements (e.g. oncogenes) on ecDNA result in amplification without the need for repeated breakpoint use and duplication events; (e) oncogenes could be expressed in a tissue specific manner providing selective advantage to different tumor sub-types. Therefore, amplicons sampled from specific tumor sub-types could be enriched in specific oncogenes, while being structurally quite different; finally, (f) ecDNA can also reintegrate into non-native chromosomal locations as homogenously staining regions (HSRs), while maintaining their structure across cell passages.

To test the tenets of this model, and get statistically more meaningful numbers, we explored the 135 AA amplicons in sample set 1 as well as 12,162 somatically amplified intervals in 2513 TCGA^23^ samples determined by CNV arrays (Methods 8). Importantly, intervals from the AA amplicons showed a significant overlap with the TCGA intervals (Methods 9, p-value = 1.1 · 10^−8^). Consistent with tenet (a), the TCGA interval size and copy number both followed exponential distributions with mean 1.74Mbp and 3.16 copies respectively (Methods 10; Fig2E, Supplementary Fig 5). Individual intervals within multi-interval AA amplicons were similar in size to single-interval amplicons; however, (see tenet (b)) complete AA amplicons containing multiple intervals from a single chromosome (14/135) (Fig 2B) or from multiple chromosomes (17/135) (Fig. 2C) were on average larger in size than the amplicons containing only one interval (101/135) (Fig2A) (Fig 2F, Methods 11, figshare). In support of tenets (c), (d), we had previously shown an increase in the copy number heterogeneity as well as an enrichment of oncogenes in ecDNA^9^.

Corroborating tenet (e), that ecDNA formation drives tumor growth through the amplifications of oncogenes which specifically confer a high selective advantage to the specific tumor sub-type, we found that amplifications of 59 distinct oncogenes were specifically enriched in 19 of 33 cancer types in the TCGA sample set (Supplementary Fig 6, Methods 12). For example, MDM4 and EGFR were enriched in glioblastoma, MYC and ERBB2 were enriched in breast cancer whereas MDM2 was enriched in both. In turn, a significant portion of the enriched oncogenes manifested in the amplicons of the corresponding cancer types from sample set 1. Limiting the analysis to the 48 enriched oncogenes in 10 TCGA sub-types that were present in sample set 1, we found that amplicons in 4 cancer types contained 18/48 oncogenes which were enriched in the corresponding TCGA cancer types while in 4 more cancer types, the corresponding TCGA samples did not show any enriched oncogenes (Supplementary Table 4).In support of tenet (f), we found no separation between intra-chromosomal (HSR) and extrachromosomal (ecDNA) amplicons in terms of their size and copy number. Detailed AA reconstructions showed that amplicons preserved their structures within biological replicates but evolved over time, in response to drug treatment, and in transition from ecDNA to HSR and back^9^ (Fig 2D, Supplementary Fig 7).

To test whether neighboring chromosomal features or functional elements outside the oncogenes played a role in amplicon formation or tumor growth in multiple samples, we measured the size of the overlap between amplified intervals containing the top 3 oncogenes - EGFR, MYC and ERBB2. For each oncogene, and each pair of samples focally amplifying the oncogene, we measured the size of the overlap to evaluate the hypothesis that the size of the overlap was significantly larger than in a null model in which overlaps are obtained from a random choice of breakpoints around the oncogene. The QQ plots of pairwise similarity scores indicated that the null hypothesis could not be rejected (Supplementary Fig 8, Methods 13). Thus, we found no evidence to suggest that recurrent breakpoint use is important for amplicon formation. These results complement previous results^2^ which also reported a lack of association with fragile sites, segmental duplications or repetitive elements in regions of complex genomic rearrangements, and they strengthen the case for an ecDNA based model of CN amplification.

In a second application of AA, we looked at focal amplifications near genomic viral integrations from 68 cervical cancer tumor samples^23^ (sample set 2) with matched normal blood samples (Methods 14, Supplementary Table 5). AA detected human papillomavirus (HPV) genomic sequence in 67/68 tumor samples and none in matched normal. We found HPV integrated into the human genome in 18/20 high-coverage(>30×) and 33/48 low-coverage(<10×) samples. AA reported that viral integrations induced formation of 41 human-viral fusion amplicons containing both viral DNA and segments from the human genome in 49% (33) of all samples (Fig 3A, Supplementary Fig 9, Supplementary Table 6, figshare). While six fusion amplicons contained an oncogene (3 with MYC, 1 each with ERBB2, BIRC3 and RAD51L1), the majority of fusion amplicons contained human sequence from intergenic regions.

**Figure 3.**
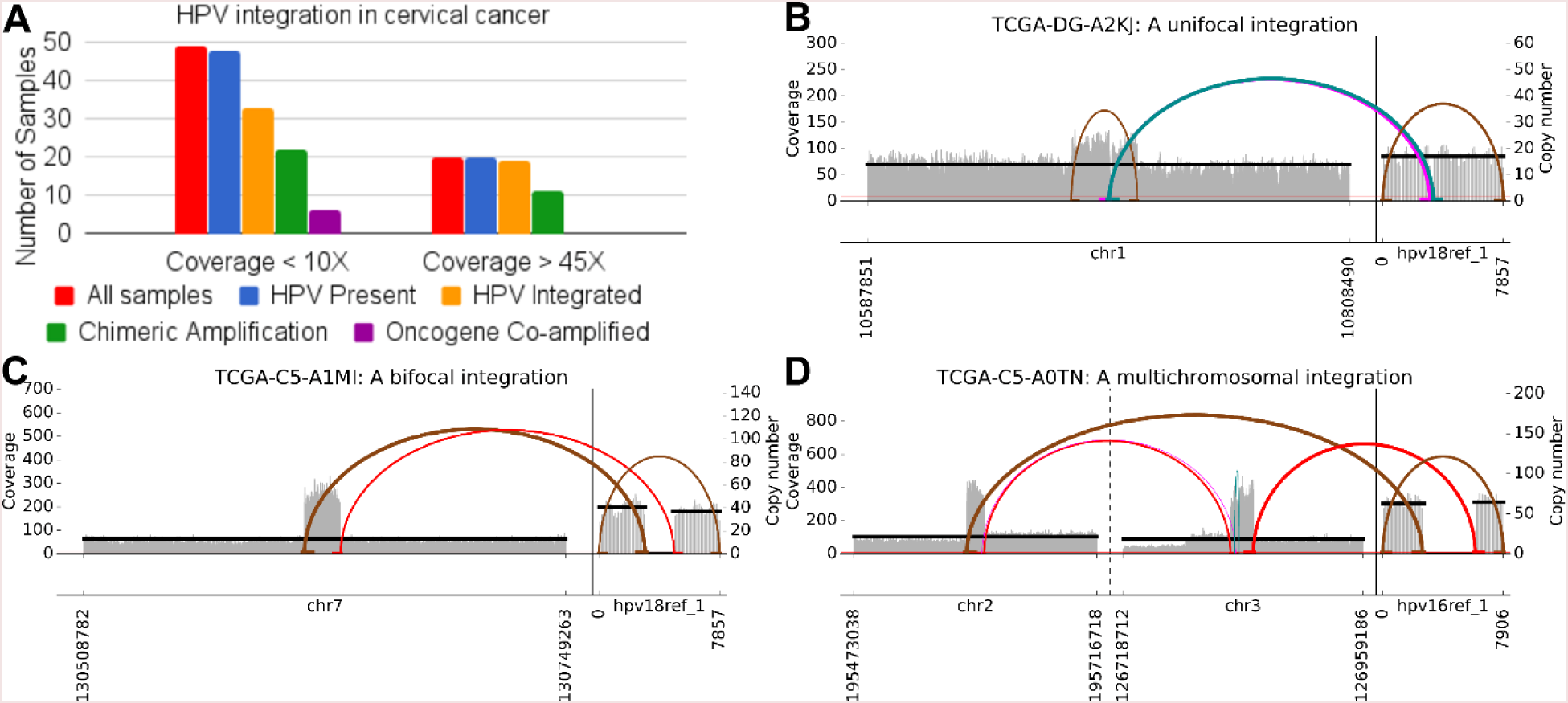
Amplicon structures near viral insertions. A) In genome sequences derived from 67 cervical cancer samples with matched normal blood, HPV sequence was identified in all 67 tumor samples (with genomic integration in 51) compared to none of matched normal blood samples. 41 fusion amplicons were reconstructed in 33 samples. B) While 14 of the viral insertions gave a unifocal amplification signature, consistent with viral insertion at a specific genomic location, (C) 32 amplicons showed a bifocal signature, (D) A 2-way bifocal signature in sample TCGA-C5-A0TN with 2 segments from chr2 and chr3 connected to a viral segment in a circular or tandemly duplicated structure with 10 copies. The prevalence of bifocal signatures is suggestive of hybrid ecDNA elements containing virus and human sequence.

The simplest mechanism of viral integration, which we will call a *unifocal integration*, consists of the virus inserting itself into the human genome by causing exactly one doublestranded break (Fig 3B). AA reconstruction revealed a novel *bifocal signature* where the endpoints of the amplified human interval were flanked by the virus (Fig 3C). For example, if 4 ordered segments ABCD represent a section of the normal human genome and V represents a viral segment, then a fusion amplicon induced by a unifocal integration might result in a structure of the form A[BVC]^n^D. In contrast, we see structures of the form AB[VB]^n^C, reminiscent of a bifocal insertion. A simpler explanation for bifocal signature is a circular extrachromosomal amplification of the form (BV) where V is connected back to B. Only 14(34%) fusion amplicons displayed a unifocal signature. In contrast, 19(46%) amplicons displayed a bifocal signature. An additional 12(29%) amplicons showed a ‘weak’ bifocal signature where only the highest copy segment was flanked by the virus but the virus did not flank neighboring amplified segments with smaller copy numbers (Supplementary Fig 9, Methods 15). Thirteen amplicons contained multiple human-viral connections. Sample TCGA-C5-A0TN contained an unusual amplicon with a 2-way bifocal signature where, 2 segments from chr2 and chr3 were connected together and the virus in turn flanked the outer end of each segment in a circular or tandemly duplicated structure with 10 copies (Fig 3D). Through simulations, we concluded that a unifocal integration followed by random rearrangements is unlikely to result in the formation of an amplicon with a bifocal signature (Methods 16, Supplementary Fig 10). Akagi et al have proposed a looping model where origins of replication on the human genome drive amplification^24^. However, the prevalence of bifocal signatures and multiple chromosomes as part of an amplicon and the ubiquitous presence of the HPV origin of replication alongside the viral oncogenes E5 and E6 in reconstructed amplicons suggest an alternative possibility that the chimeric amplification could be mediated through ecDNA formation. Although episomal virus in its native form has been reported extensively in cancer cells, the AA reconstructions make a compelling case to investigate the presence of fusion human viral segments in the form of ecDNA.

AA is a robust and viable tool for reconstructing possible ecDNA and other focal amplicon structures from short-read data and allows for an interactive exploration of alternative structures. This method and subsequent analysis on a pan-cancer data set suggests that formation of ecDNA could play an important role in creating the complex rearrangements and copy number increases across the spectrum of cancer subtypes.

## Supplemental figures

- Fig S1: Example of the cycle merging operation
- Fig S2: Error model for benchmarking AA
- Fig S3: Accuracy of AA submodules and reconstruction
- Fig S4: Runtime of AA
- Fig S5: Distribution of amplicon intervals corresponding to Fig 2E
- Fig S6: Cancer type-specific oncogene enrichment
- Fig S7: AA amplicons in biological replicates for 7 cancer samples evolving over the passage of time or in response to drug treatment
- Fig S8: QQ-plot of similarity of overlapping amplicon intervals vs expected similarity
- Fig S9: Size and copy number distribution of human-viral fusion amplicons in cervical cancer
- Fig S10: Unifocal and bifocal signatures over evolution of fusion amplicons

## Supplementary tables

1. Sample list
2. Seed intervals
3. Final amplicon intervals, oncogenes and classification (single or multi-interval)
4. Oncogene amplified in corresponding types in TCGA and sample set 1
5. Viral sample list with detected HPV strain and integrations
6. List of viral amplicons, oncogenes and classification (unifocal/bifocal)

## Figshare

HYPERLINK “https://figshare.com/articles/AmpliconArchitect_reconstructions/5950339“ https://figshare.com/articles/AmpliconArchitectreconstructions/5950339

- All reconstructions on previously reported amplicons
- All reconstructions from sample set 1 and replicates
- All examples of simple cycles
- All multi-interval amplicons
- All multi-chromosomal amplicons
- Sample with Breakage-Fusion-Bridge signature

## Code and data availability

The AmpliconArchitect software described in the manuscript is available at https://github.com/virajbdeshpande/AmpliconArchitect. Whole-genome sequencing data for sample set 1 and 6 replicates for sample GBM39 were downloaded from the NCBI Sequence Read Archive (SRA) under Bioproject (accession number: PRJNA338012). 12 replicates for other samples are available on SRA under Bioproject (accession number: https://www.ncbi.nlm.nih.gov/bioproject/PRJNA437014). The reconstructions described in this manuscript may be downloaded from https://figshare.com/articles/AmpliconArchitectreconstructions/5950339

## Methods

### 1 Seed interval selection: Interval merging and copy number threshold

AA requires a seed interval in addition to mapped genomic reads. The seed interval serves as a starting point for AA to search for all connected genomic intervals contained in the amplicon. Here, we pick seed intervals in two different sample sets: a) WGS of tumor samples across multiple cancer types, b) WGS of cervical cancer samples infected with HPV.

A) To identify the set of seed intervals in the WGS samples from the pan-cancer dataset, we defined parameters CN_THRESHOLD and SIZE_THRESHOLD for minimum bounds on copy number and size of the interval. Aiming to find a criterion for identifying somatically amplified intervals, we compared the CNV calls for matched tumor and normal samples downloaded from TCGA consisting of 22376 masked CNV call files from TCGA generated from Affymetrix 6.0 data for 10995 matched cases. For a given CN_THRESHOLD, define an *amplified segment* as a single CNV call with copy number greater than CN_THRESHOLD. A sample might have multiple amplified segments adjacent to each other. The size of the amplified segment was simply the number of base-pairs in the segment. For each sample, we merged consecutive amplified segments within 300kbp of each other to create the set of *amplified intervals.* The size of amplified interval was the sum of sizes of the amplified segments in the interval and the copy number of the amplified interval as the weighted average of the amplified segments weighted by their sizes. We counted the number of amplified intervals in all normal and tumor samples in the TCGA set for values of CN_THRESHOLD in {3, 4, 5, 7, 10} and SIZE_THRESHOLD in {10kbp, 50kbp, 100kbp, 200kbp and 500kbp}. Based on these, we chose the values CN_THRESHOLD=5 and SIZE_THRESHOLD=100kbp for selecting the seed intervals which resulted in 25 intervals in normal samples and 12162 intervals in tumor samples which did not have a corresponding amplification in the normal samples.

Next, we analyzed the WGS reads from the WGS samples from the pan-cancer dataset. We mapped the reads to the hg19 genome from UCSC genome browser^25,26^ and obtained CNV calls using the CNV calling tool ReadDepth CNV software version 0.9.8.4^22^ with parameters FDR= 0.05 and overDispersion parameter=1. We also obtained the CNV calls for 8 normal control samples and created the set of amplified intervals from the CNV call sets for all samples with CN_THRESHOLD=5 and SIZE_THRESHOLD=100kbp. First, we looked at the amplified intervals from the normal samples and found genomic regions which were amplified in 2 or more normal samples. These regions were marked as *blacklisted regions.* Since ReadDepth did not report CNV calls for chrX and chrY, we used a previously computed list of recurrent CNVs on the X and Y chromosome reported by Layer et al^13^.

From the amplified intervals from WGS of tumor samples, we filtered out false intervals using 3 criteria:

(i) We identified intervals overlapping blacklisted regions and trimmed them to exclude the portions within 1Mbp of the blacklist regions.

(ii) For each interval, we calculated its average repetitiveness by defining Duke35 repetitiveness score based on the mappability score track from UCSC genome browser^26^. This mappability score track reports the repeat count of each 35 bp window in the reference genome up to copy number 5. We computed the Duke35 repetitiveness score of an interval as the average score of all 35bp windows in the interval and filtered out intervals with score > 2.5.

(iii) We looked at intervals overlapping regions of segmental duplication (SD) reported by the human paralog project. For these intervals, we defined an *SD-adjusted copy number* as the interval copy number downscaled by its average repeat count. In the absence of information regarding actual repeat counts of the SDs, we assumed a repeat count of 2. As a result, the SD- adjusted CN = Interval CN / (1 + Total length of overlapping SDs / length of interval). We only retained amplified intervals if their SD-adjusted CN was great than 5.

Finally, to correct for the copy number gain due caused due to aneuploidy, we required that the difference in copy number of the interval and the median copy number of the chromosomal arm should be at least 3. Specifically, for amplicons in chromosomes with reference copy number of two, the copy number cutoff was 2 + 3 = 5 = CN_THRESHOLD. We applied these filters on the ReadDepth calls for the 117 WGS tumor samples to obtain 255 intervals in 55 samples.

B) For detecting chimeric human-viral amplification in cervical cancer samples, we created a combined reference genome consisting of the human chromosomes and the viral reference genome and aligned reads to this combined reference (Methods 13). We selected the viral genome as the seed interval. This was a highly selective criterion for selection of the seed interval which allowed us to perform a more sensitive search for amplicons. As a result, there was no initial cutoff on copy number or size of the seed interval. The cutoffs were chosen at the end of final reconstruction.

### 2) AA methods

The AA pipeline starts with the seed interval and mapped reads and performs multiple steps including search for amplicon intervals, detection of genomic rearrangements, construction of breakpoint graph, decomposition of the graph into simple cycles and visualization of the cycles. To perform these steps efficiently and accurately, AA implements and uses multiple low-level modules. Here, we briefly describe the implementation of AA pipeline and the low-level modules. The AA software described here may be downloaded from https://github.com/virajbdeshpande/AmpliconArchitect commit id: d993372.

#### A) Low level modules

##### (i) Sequencing parameter estimation

This module estimated the parameters of sequencing coverage and variability as a function of window size, as well as the read and fragment insert lengths of the sequencing library. Given a bam file of mapped reads and a window size ws, AA obtained an initial estimate of the median coverage for 1000 randomly chosen windows from non-blacklisted regions, excluded all windows with coverage = 0 or > 5 times the initial median and recalculated the mean μ_ws_), median (*θ*_WS_) and standard deviation (*σ*_ws_) of window coverages for the given window size. AA computed the coverage for window sizes ws=10kbp and ws=300bp. Further, it obtained the read pairs from the windows used for estimating the coverage with window-size 10kbp, estimated the fraction P of “properly mapping” read pairs and estimated the mean read length R and the mean (μ(l)) and standard deviation (σ(l)) of the fragment insert length of the properly mapping read pairs as reported by the read aligner in the SAM alignment flags.

##### (ii) CNV boundary detection

This module identified positions of copy number changes in an interval using only the coverage histogram. It first identified an initial list of boundaries of CNV segments based on a histogram with window size 10kbp and then refined these boundaries through a local search by based on a histogram with window size 300bp within the neighborhood of the initial list of boundaries. To estimate the CNV boundaries for a given interval size, AA used a meanshift procedure adapted from Abyzov et al^20^. In the meanshift procedure, AA identified the CNV boundaries as the locations of the minima of the Gaussian kernel density estimator indicative of a large change in coverage. Specifically, for each window “i”, define a Gaussian kernel density function *F_i_*

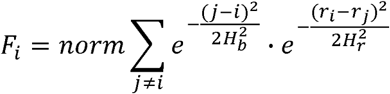

Here j iterates over 50 neighboring windows on either side of window i, r_i_, r_j_ are the coverage depths for bins i and j, H_b_ is the bandwidth for bin index and H_r_ is the bandwidth for the coverage depth. For bin i with size ws and coverage c_i_, AA set 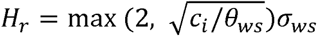. H_b_ iteratively took values in the order (2, 5, 10, 50, 100) as described below. *norm* represents the constant normalization coefficient. Thus, (VF)i, the component of the gradient of F_É_ along the genomic coordinates is:

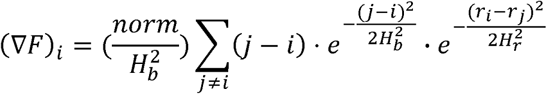

AA selected the boundaries between pairs of consecutive windows where (VFchanges from negative to positive, merged all windows between these boundaries into segments and calculated the average coverage C_s_ for each segment s. AA selected the boundaries where the difference in coverage |C_s1_ - C_s2_| for consecutive segments s1 and s2, was found to be significant as described below. If either segment was smaller than 15 windows, then it required |C_s1_ - C_s2_| > 3ct_ws_- max(C_s1_, C_s2_) / 0_WS_. AA detected the meanshift boundaries at various scales by iteratively increasing the size bandwidth from 2 to 100 windows while freezing segment boundaries it found significant in each stage. AA obtained an initial high confidence set of CNV boundaries in an interval by searching for the meanshift boundaries in the entire interval with window size ws=10kbp. It then refined these boundaries by running the meanshift algorithm with window size ws=300bp and from the new call set, picking the new boundary with desired directionality change in coverage and largest difference in coverage of adjacent segments.

The CNV boundary detection module calculates the average coverage for genomic segments and defines the *coverage ratio* as *CRs =* 2*C_s_* / μ_*ws*_. The module could be run in two modes. In the sensitive mode, the difference in coverage of adjacent segments |C_s1_ - C_s2_| was considered significant as determined by independent t-test of the distribution of window coverage of s1 and s2. In the default mode, the module further filtered out boundaries if the difference in coverage ratios 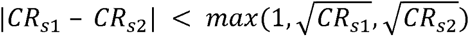. The sensitive mode was specifically used for chimeric human-viral amplicons where the virus could have very high copy number as compared to the human intervals due to independent amplification. The default mode was used for all other amplicons in order to focus on amplicon structures that comparable in abundance with the highest copy structure and ignore noise from low copy rearrangements.

##### (iii) Breakpoint detection

This module took as input one or more intervals, and identified all breakpoints associated with these intervals using discordantly mapping read pairs:

a. Identify all discordant reads mapping to the intervals such that the mate maps either maps to a different chromosome, has unexpected mapping orientation, or maps at location such that the distance between the outermost mapping positions of the read pairs is outside of the range (μ(!) - 3σ(l), μ(l) + 3σ(l)). Cluster reads that map within (μ(l) + 3σ(l)) basepairs of each other and have the same mapping orientation.
b. For reads with each cluster identify mapping positions of the mates and create one or more cluster pairs, “biclusters” from read pairs (including secondary alignments) where the first cluster consists of subset of reads from cluster from step(a) and that the second cluster corresponds to mates of these reads that map within (μ(l) + 3σ(l)) of each other.
c. For each bicluster, filter reads in repetitive regions with MAPQ ≤ 5 or satisfying one of the 3 criteria for filtering out repetitive intervals described in the seed interval selection process.
d. Remove biclusters whose size is smaller than a significance threshold S as described below. The size of a bicluster is the number of unique read pairs in the bicluster.
e. Report pairs of breakpoint inferred from bicluster.

The significance threshold for number of read pairs in the bicluster could be chosen from 4 different options used for specific scenarios: (a) a fixed input parameter (e.g., 2 read pairs for a sensitive search), (b) minimum number of read pairs determined by the average sequencing coverage, read length and fragment insert length, (c) minimum number of read pairs for a region with copy number estimated by the coverage ratio of the meanshift segments, or (d) minimum number of read pairs determined by the difference in the coverage ratio across the CNV boundary. The minimum number of read pairs S associated with a given CR or difference in CR of segments was calculated as *S_CR_* = *P* × *CR* × *μ*_300_ ×(*I* − *R*) / 2R / D. Here D = 20 was a downscaling factor which we chose based on observations from our simulations which suggested that the expected number of read pairs scaled down 20 times provided a classifier with high sensitivity without affecting specificity at multiple copy number states. Selection and hard-coding of the parameter D was the only fingerprint of “training” AA based on the simulation “evaluation set”, otherwise all development of AA was done prior to evaluation on the simulated examples.

#### B) AA pipeline

AA implemented a series of steps to start from a seed interval and ultimately reconstruct the full structure of the amplicon and provided informative results from each stage:

##### (i) Interval search

In this step, AA started with the seed interval and iteratively identified the list of intervals belonging to the amplicon. It started by creating a max-heap data-structure storing the seed interval.

a. AA repeated the following steps until the max-heap was empty or after 10 iterations (Fig 1Aii).
  i. In each iteration, AA selected an interval and determined the discordant read-pair biclusters in a CN-sensitive fashion and selected biclusters with the mate mapping outside previously seen intervals.
  ii. It then attempted to extend the bicluster by querying whether the extended portion is amplified. A query segment was classified as *amplified* if it had at least 20% of windows(ws=10kbp) with coverage > *θ* _10000_+ 3σ_10000_, or if it was smaller than 20kbp and contained at least 2 discordant edges (bicluster size corresponding to CN=2). AA then efficiently extended the query bicluster by iteratively doubling the size of the extended portion until the extension is found to be amplified and then iteratively reduced the extension query size by half. If AA was able to successfully extend the query bicluster, then it extended it further by 100kbp and recorded the extended interval for future iterations.
  iii. After AA recorded all amplified neighbors, AA marked the interval as *seen*, updated the max-heap ordered by the number of discordant read pairs connected to previously seen intervals and greedily picked the interval at the top of the heap for the next iteration.
b. AA reported all amplified intervals from the extension step.

##### (ii) Interval rearrangements, partition and visualization

a. AA calculated all the coverage meanshift boundaries and initial copy number estimates for corresponding segments (Fig 1Aiii).
b. It then created the list of discordant read pair biclusters with bicluster size thresholds determined by the CN estimates (or differences) (Fig 1Aiv).
c. Additionally, for meanshift boundaries where it did not find a matching discordant read biclusters, it performed a sensitive local search for discordant read pairs with a bicluster size threshold of just 2 read pairs (Fig 1Av).
d. Finally, it created the set of genomic locations of all rearrangements with a discordant read pair bicluster or a meanshift boundaries with no matching discordant reads. Using this set of locations, it partitioned all the intervals into *sequence edges.*
e. For the output from the second stage, AA created a single plot called the *SVview* which displayed the interval set with the coverage histogram, initial copy number estimates of the meanshift segments and the discordant read biclusters.

##### (iii) Breakpoint graph and copy number estimation

AA used the sequence edge partitions to construct a breakpoint graph^27^.

a. For each sequence edge, it created 2 breakpoint vertices marking the start and end of the genomic segment and added a sequence edge connecting the 2 vertices.
b. AA augmented the vertex set with a special *source vertex.*
c. For each discordant read bicluster, it added a *discordant breakpoint edge* connecting the respective endpoints of the corresponding sequence edges if both the clusters belonged to the interval set.
d. For all biclusters with one cluster outside the interval set, it introduced a *source breakpoint edge* connecting the source vertex to the breakpoint vertices corresponding to the clusters within the amplicon intervals.
e. It also added *source breakpoint edges* connecting the source vertex to breakpoint vertices corresponding to meanshift vertices with no corresponding discordant biclusters and to end points of the amplicon intervals.
f. Finally, AA connected breakpoint pairs corresponding to consecutive sequence edges within each interval with *concordant breakpoint edges*.
g. For each sequence edge and breakpoint edge, AA recorded the number of reads and read pairs respectively mapping to the edge.

A correctly reconstructed breakpoint graph represents a superimposition of all amplicon structures. Each cyclic structure forms a cycle of alternating sequence and breakpoint edges. A linear structure with end-points connected to genomic positions outside the cycle, can be represented as an alternating closed walk starting and ending at the source vertex. The breakpoint graph construction may not always be complete due to missing edges leading to inaccuracies in final prediction of the structures. This problem was partially alleviated by the fact that, if AA failed to detect a discordant breakpoint edge, but detected the locations of the rearrangement through meanshift boundaries, then the corresponding amplicon structures were represented as one or more walks starting and ending at the source vertex. Henceforth, we restrict the definition of *cycle* in the breakpoint graph, as a closed walk with alternating sequence edges and breakpoint edges with the exception that the walk may contain 2 consecutive breakpoint edges connected to the source at most once. Next, we note that the CN of each edge is the sum of CNs of each amplicon structure where each traversal of the edge separately. As a result, the copy numbers in the graph follow a balanced flow property wherein the CN of a sequence edge matches the sums CNs of breakpoint edges connected to each breakpoint vertex of the sequence edge. AA modelled the number of read fragments mapping to each edge as a Poisson distribution with the parameters determined by the CN, edge length and sequence coverage parameters. Under this model, AA estimated the copy number CN_seq_ for each sequence edge and CN_bp_ for each breakpoint edge by optimizing a balanced flow (linear constraints) with the convex objective function^21^:

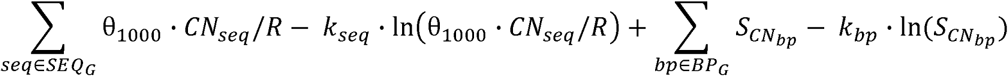

where SEQ_G_ represents all sequence edges and BP_G_ represents all breakpoint edges in breakpoint graph G, k_seq_ and k_bp_ represent the number of reads mapping to sequence edge seq and the number of read pairs mapping across breakpoint edge bp respectively with the constraint:

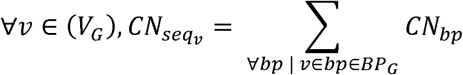

where seq_v_ represents the sequence edge connected to breakpoint vertex v, V_G_ represents set of breakpoint vertices in breakpoint graph G. The optimal solution for the balanced flow was obtained using the convex optimization package Mosek version 8.0.0.60^28^. For the third stage output, AA reported the graph edges and their copy counts as text output.

##### (iv) Cycle decomposition

As described above, a linear or cyclic amplicon structure can be represented as one or more cycles in the breakpoint graph. However, even with correct reconstruction and CN assignment to the breakpoint graph, the cycles cannot always be inferred unambiguously, especially with repeated traversals of an edge. Conversely, there may be two or more possible sets of cycles and associated copy numbers, such that combining the cycles within each set may finally result in the same breakpoint graph with the same copy number assignments. Here *combination of cycles* simply means summing up the copy numbers for each graph edge from each cycle. It is not always practical to enumerate all possible amplicon structures because the number of possible structures can be exponentially large. To address this issue, we first observed that a cycle traversing an edge multiple times in the same direction can be divided into two smaller cycles and conversely, the two cycles can be merged to form the original cycle. However, if we iteratively merge multiple cycles together, then changing the order in which the cycles are merged can produce different resulting structures all with the same edges and copy counts. Based on this observation, AA decomposed the breakpoint graph in to *simple cycles*, with the aim to represent a large number of amplicon structures using relatively few cycles. We defined a *simple cycle* as a cycle which traverses any sequence edge at most once in each direction and hence cannot be divided into smaller cycles. We defined the *decomposition of the breakpoint graph* as a set of simple cycles with CN assignment such that the CNs of any edges in all the simple cycles sum up to the CN in the breakpoint graph. While a breakpoint graph may have multiple decompositions and the simple cycles in a single decomposition may not be always be combined to form every possible amplicon structure, these cases require the breakpoint graph to have certain complex patterns expected to occur in a small fraction of amplicons. Instead, AA decomposed the breakpoint graph using a polynomial time heuristic which iteratively picked the simple cycle with the highest CN and decremented the CN from the corresponding edges in the breakpoint graph. With this algorithm, AA could prioritize the structures that had the highest CN as well as the cycles which occur in a large number of structures providing a meaningful way of highlighting the important features of the amplicon. AA provided a text file containing the ordered list of segments within in each simple cycle. Additionally, AA provided an interactive visualization of the simple cycles called the “CycleView” which could be merged to into larger cycles to investigate possible amplicon structures. In the CycleView, AA displayed the segments aligned with their genomic position in the SVview and consecutive segments were placed on consecutive rows. If two cycles contained overlapping segments, then a user could select the cycles, their overlapping segments and merge the cycles to form larger cycles. The CycleView provided a way to interpret the structure of the cycle while visualizing the genomic location and annotations.

#### 3) Samples reported by other studies

We ran AA on previously reported amplicon and provide a comparison of AA reconstructions with previous studies (Supplementary Materials). The samples included:

Dataset (i) contained 6 samples (HL-60, GLC-1-DM, GLC-2, GLC-3, COLO320-DM, COLO320HSR) provided to us by the authors of the original paper. Each sample was predicted by the original study to contain an amplicon with the oncogene MYC, along with PCR validation of breakpoint edges. We mapped the WGS samples to with with coverage between 4.6X to 10.5X and remapped the reads to hg19 reference genome with BWA MEM. We picked the seed intervals using the ReadDepth as described on Online methods section 1.

Dataset (ii) had 3 glioblastoma samples (TCGA-06-0648, TCGA-06-0145, TCGA-06- 0152) from Sanborn et al^19^, also studied by Dzamba et al^18^. 1 ovarian cancer sample from Oesper et al^15^ (TCGA-13-0723) also studied by Dzamba et al^18^. We downsampled the bam files to coverage between 4X-7X by selecting read pairs with specific read group identifiers. The read group identifiers were selected to be sets of identifiers with the same read length and roughly similar insert length. The exact identifiers selected are mentioned in Supplemental materials. We picked seed intervals based on calls from CNV calling tool ReadDepth with copy number > 5 and size > 100kbp as described in Methods 1A.

Dataset (iii) consisted of 12 HPV infected cancer samples (HNSCC and CESC) from Akagi et a^24^. For each sample we predicted the HPV strain as described in Methods 13, remapped all the reads to the combined reference by concatenating the hg19 human reference genome to the reference genome of predicted HPV strain, and used the interval corresponding to the predicted HPV genome as the seed interval.

#### 4) Simulation algorithm

We developed a simulation algorithm, AAsim, to simulate 960 amplicons with known “true” structures to measure the accuracy of AA. Simulations of AAsim can be flexibly adjusted of multiple input parameters including: i) Interval size, ii) Copy number, iii) Number of rearrangements, iv) Probability of duplication and v) Depth of coverage. To allow testing the reconstruction without the bias of seed selection, AAsim simulated viral (HPV16) human hybrid structures. The HPV16 genome served as a de facto seed interval such that AA could be tested without providing any additional information about the span of the amplicon. AAsim simulated ecDNA structures through the following steps:

1. Chose a random location on the human genome and integrate the HPV16 genome at this location.
2. Randomly select an interval of input interval size around the site of integration and circularize it to create an ecDNA element containing the human interval with the integrated virus.
3. Iteratively perform rearrangements on the ecDNA, including deletions, duplications, inversions and translocations such that in each iteration. Each type of rearrangement is selected with preset probabilities, as described below. The breakpoint coordinates for each rearrangement are chosen uniformly at random from the entire ecDNA structure, but rearrangements that deleted the viral genome entirely were not permitted. Iterations were performed until the required number of rearrangements were induced.
4. Record the order of segments in the final structure to be used as “truth” set and assigned a copy count to the ecDNA using the input parameter.
5. Generate 100bp paired-end reads from the ecDNA using the ART Illumina read simulator^29^ with given depth of coverage. Reads are also generated from other regions of the reference genome for AA to estimate the profile of the sequencing depth.

The output of AA sim included the target “true” amplicon structures and the paired-end reads simulated for these amplicons. The values chosen for the input parameters were: i) Interval size: 40kbp, 160kbp, 640kbp, 2.4Mbp; ii) Copy number: 4,16, 32; iii) Number of rearrangements: 0, 4,8, 16, 32; iv) Probability of duplication: 0, 0.25, 0.5, 0.75; v) Depth of coverage: 1, 4, 16, 32. In total, we generated 960 simulations for all combinations of these parameters. The sets of simulations with increasing duplication probability presented test sets with increased difficulty of reconstruction due to larger number of cycles per structures as well as larger number of segments. In order to estimate the runtime of AA for larger amplicons, we simulated 24 more structures with parameters: i) Interval size: 5Mbp, 10Mbp; ii) Copy number: 32; iii) Number of rearrangements: 0, 8, 32; iv) Probability of duplication: 0, 0.25, 0.5, 0.75; v) Depth of coverage: 32.

#### 5a) Accuracy of methods for SV and CNV analysis

SV and CNV analyses provide the building blocks for AA to accurately reconstruct the breakpoint graph and ultimately predict the full structure. In order to establish the reliability of these critical components, we measured the accuracy of methods for interval detection, discordant edge detection and meanshift edge detection. In terms of interval detection, even though AAsim simulated an ecDNA circularized from a single interval, the final structure can have multiple intervals due to deletion of intermediate segments. We measured the number of intervals completely identified by AA (TP), the number of intervals not completely identified by AA (FN) as well as the number intervals reported by AA which did not have any amplification (FP). For measuring the accuracy of CNV detection, we measured the number of CNV boundaries correctly (within 10kbp) identified by the meanshift edge detection algorithm (TP), the number of CNV boundaries not identified (FN), as well as the number of locations reported which actually did not have a copy number change (FP). Similarly, for discordant edges, we reported the number of edges with both breakpoints predicted correctly (within 300bp) (TP), one or more breakpoints not detected (FN) and discordant edges reported which did not exist in the true structure (FP) (Supplementary Fig 3).

#### 5b) Edit distance computation and reporting

For measuring the accuracy of final amplicon reconstruction, we defined a distance measure to quantify the difference between the predicted structure as compared to the true structure from the simulation. Inspired by the genome sorting problem, the goal of the distance measure is to represent the number of operations to transform the predicted structure into the true structure. The genome sorting problem aims to find the distance between two related genomes by counting the minimum number of operations to transform one genome into another. These operations may include inversions^27^, translocations^30^ or double-cut-and-join(DCJ) operations^31^. We adapted the DCJ operation to measure reconstruction accuracy by defining the Repeat-DCJ (RDCJ) distance which can separately count the portion of operations caused due to reconstruction errors and due to alternative traversals across repeats. See Supplementary Figure 2. The RDCJ distance is defined as the sum of a two-part measure: i) *repeat branch swaps* and ii) *reconstruction errors.* First, the prediction may have errors caused by inaccurate breakpoint graph construction including missing or false breakpoint edges and inaccurate copy numbers of segments. We denote these as reconstruction errors as they represent errors caused by the reconstruction algorithm. Reconstruction errors involve addition and deletion of breakpoint edges. On the other hand, under perfect graph construction, the only operation needed to transform the predicted structure into the true structure is to transform the order of traversal across repeated segments without any change in the set of breakpoint edges used. We call this operation a repeat branch swap. Two cycles can be merged together through a single repeat branch swap. In measuring the RDCJ-distance, we simultaneously minimize the number of reconstruction error corrections and repeat branch swaps required for the transformation.

Under the RDCJ model, we count the edits on each segment independently and sum these up to obtain the total distance for the entire reconstruction. To achieve this, we represent each segment by a switch. A *switch* is a bipartite graph where the two parts represent the start and end of the segment respectively and the vertices are the union of breakpoint vertices from the true and predicted structures. If a breakpoint vertex is traversed multiple times in either of the structures, then we create multiple copies of the vertex equal to the maximum number of traversals in the two structures. We define a *switch graph* as a graph which consists of all the switches as subgraphs connected through *connective edges* which are union of the breakpoint edges from the breakpoint graphs of the true and predicted structures with appropriate multiplicities. Each switch vertex has exactly 1 connective edge. Finally, each of the true and predicted structures form a walk on the switch graph inducing respective matchings within the bipartite switches. The edges within the matching, called *match edges*, connect consecutive breakpoint edges in the structure. As a result, we represent the edit distance of the predicted structure to the true structure by counting the number of operations of transforming each switch independently.

Consider a switch with vertices V_1_, V_3_ on one shore, and V_2_, V_4_ on the other shore of the bipartite graph with switch edges (V_1_, V_2_) and (V_3_, V_4_). In using a repeat branch swap to transform the predicted matching to the true one, we can, for example, replace edges (V_1_, V_2_) and (V_3_, V_4_) by 2 new edges (V_1_, V_4_) and (V_3_, V_2_). Note that a repeat branch swap represents a copy number neutral operation and correspondingly, the number of switch edges does not change. Since the set of vertices in the matching is invariant, the set of connective edges connected to matched vertices also does not change. In contrast, correction of reconstruction errors involves addition, deletion or reassignment of exactly 1 vertex of a match edge.

To measure the accuracy of AA, we counted and reported the required the number of the operations of both types across all switches for each simulated structure. We classified the 960 simulated structures on 4 groups based on the value of the parameter probability of duplication, representing amplicons which are increasingly difficult to reconstruct. To provide a standard yardstick, we compared the reconstruction errors of AA predictions against those of a random predictor. The random amplicon structure consisted of a randomly shuffled order of all segments from the true amplicon structure. Notably, the average performance of the random predictor closely followed 2 times number of segments in the amplicon where number of segments is measured by adding up counts of segments. Based on this observation, we defined the *error rate* of one or more predictions as the total reconstruction errors as a percentage of 2 x total number of segments (Fig 1h-k, Supplementary Fig 3).

### 6) Runtime computation

We recorded the runtime of AA for each simulated amplicon using the Python function time.time(). We plotted a scatterplot of the runtime as a function of the total DNA content of the amplicon on a logscale graph to capture performance on small and large amplicons (Supplementary Fig 4). The total DNA content was defined as length of ecDNA structure X copy number X depth of coverage. We plotted the best fit line on the logscale graph for the runtime as function of the total DNA content using the linregress function available in the Python library scipy.stats.

### 7) SRA samples, ReadDepth CNV calls

We used sequencing data from 117 cancer samples including cell lines, patient-derived xenografts (PDX) and tissue samples and 8 normal control samples originally described in Turner et al^9^. These samples may be downloaded from NCBI Sequence Read Archive (SRA) under Bioproject (accession number: PRJNA338012). Additionally, we studied WGS of 18 biological replicates of 7 samples totaling to 135 cancer WGS datasets. 4 of these samples had replicates treated with targeted drugs and glioblastoma PDX GBM39 also had post-treatment replicates (Table S1). FISH results for oncogene probes reported in Turner et al were used to mark amplicons to be present on EC only, HSR only or both EC and HSR (Fig 2D, Table 3).

After reconstruction of amplicons in 117 WGS tumor samples using the 255 seed intervals from ReadDepth, AA reconstructed 135 amplicons which consisted of 265 intervals. While AA merged multiple seed intervals into larger intervals, AA reported 63 new intervals not intersecting the seed intervals including possible false positives from repetitive regions (Table 2, 3).

### 8) TCGA interval set and somatic CNV identification

We downloaded 22376 masked CNV call files from TCGA^23^ generated from Affymetrix6. 0 data for 10995 cases. We mapped the original calls from hg38 coordinates to hg19 coordinates using the Liftover tool from UCSC genome browser. We selected 10494 cases for which Liftover successfully mapped the calls for at least 1 cancer sample and 1 matched normal sample. For cases with multiple call sets, we took the average copy number of each segment for cancer and normal call sets respectively. Next, we selected the CNV calls in the tumor samples according to the seed interval selection procedure described in Methods 1. However, we did not exclude the calls from the blacklisted regions since we believe those regions only need to be blacklisted due to artefacts specific to WGS samples and not array data. Using this method, we obtained 12162 intervals in 2527 cancer cases. Comparing the copy number calls in cancer samples with copy numbers in matched normal samples, we filtered out calls where the difference in copy number was less than 3 (specifically, copy number at least 5 when there was no CNV call in the normal sample). This criterion only filtered out a small number of intervals resulting in 12162 intervals in 2513 cancer cases. We used this set for further analysis.

### 9) Overlap of AA amplicons with intervals amplified in TCGA

We tested whether amplicons reconstructed by AA in sample set 1 were representative of the focal amplifications across human cancer by testing whether the overlap between AA amplicons with TCGA intervals was significantly larger than expected by random change. As originally described in Turner et al^9^, for each sample, we computed a match score between the AA amplicons for the sample and the TCGA intervals from the corresponding cancer types which were amplified with frequency > 1%. The match score for the sample was simply the sum of frequencies of the TCGA intervals within the corresponding cancer types that overlapped an amplicon from the sample. We recorded the cumulative match score as the sum of match scores for all samples in sample set 1.

To test if the cumulative match score for the TCGA intervals was significantly larger than expected by random chance, we generated 720 million permutations of the TCGA intervals which were amplified with frequency > 1% in each cancer type by assigning random positions to the intervals within the human reference genome while maintaining their size. We computed the cumulative match score of each permutation with sample set 1 using the same procedure as above and found that 8 permutations had a larger match score than the original TCGA intervals. The p-value of the significance of overlap between amplicons in sample set 1 and TCGA intervals was reported as 8/720 million = 1.1 · 10^−8^

### 10) Size and copy number determination and exponential distribution

The size of an amplified interval was defined as the sum of sizes of all amplified segments with copy number > 5 within the interval. The size of the amplicon was the sum of sizes of all intervals in the amplicon. The copy number of an interval was the average copy of the amplified segments in the interval weighted by their size. Similarly, the average copy number of an amplicon was the weighted average of amplified segments in all intervals of the amplicon weighted by their size. In the analysis of final reconstructions, we used the copy numbers assigned to sequence edges by AA rather than CNV calls from ReadDepth. We plotted the scatter plot for copy number vs size of the 135 AA amplicons for the 117 samples and the TCGA intervals (Fig 2d). For direct comparison, we also plotted the copy number vs size of the AA intervals (Supplementary Fig 5). We observed 6 AA amplicons and 3 AA intervals with sizes > 3Mbp and copy number > 30, whereas we observed exactly 1 TCGA interval satisfying these constraints. Particularly, the CNV array calls were capped off at copy number around 40. We verified that this was the case for the amplicons from all 4 TCGA WGS sample from previous publications where the sequencing data reported higher copy counts.

Next, we plotted histograms for the copy number and size of the TCGA intervals with bin sizes of copy number 1 and size 400kbp where the height of the histogram was log-scaled. (Fig 2d). Both histograms displayed a linear decay indicating exponential distributions. We obtained the best fit lines for these histograms using the polyfit function in the Python package Numpy based on 20 bins each for copy number (5-25) and sizes (0bp-8Mbp) beyond which the data became too sparse. Estimates for the means of the exponential distributions, 3.16 copies and 1. 74 Mbp, were obtained using the negative inverse of the slop of the best fit lines in both cases.

### 11) Determination of multi-interval and multi-chromosomal amplicons

We compared the sizes amplicons containing a single genomic interval as compared to amplicons with multiple intervals from one or more chromosomes. To be more conservative in the interval selection and avoid false intervals detected by AA from repetitive regions, we selected intervals based on segments reported to be amplified by the meanshift based CNV analysis. As a result, only high confidence intervals detected by the CNV analysis at a resolution of 10kbp were selected. Next, for a uniform definition of interval, only for this analysis, we merged all CNV segments within 5Mbp of a neighboring segment into a single interval. After merging these segments, we classified the amplicons into 3 categories: (i) Clustered: 104 amplicons containing a single interval, (ii) MultiCluster: 14 amplicons containing multiple intervals from the same chromosome and (iii) MultiChrom: 17 amplicons containing multiple intervals from multiple chromosomes. (Fig 2e).

We plotted the distribution of amplicon sizes of the entire amplicon for each of the 3 categories as well as the size distribution of the set of all intervals in all amplicons in the MultiCluster and MultiChrom categories using the Python Seaborn library function “violinplot”. We compared the sizes of all amplicons and intervals in the MultiCluster and MultiChrom category to the sizes of all Clustered amplicons using RankSum test available through Python Scipy.Stats library function “ranksums”. We observed that the MultiCluster (p=1.5810^−2^, mean=4.7Mbp) and MultiChrom (p=6.18·10^−4^, mean=7.7Mbp) amplicons were significantly larger than the Clustered amplicons (mean=2.3Mbp). However, the sizes of intervals from all amplicons in the MultiCluster (mean=2.2Mbp) and MultiChrom (mean=2.7Mbp) categories did not show any significant difference from the Clustered amplicons.

### 12) TCGA subtype specific enrichment

We considered the significance that the amplicons amplified specifically amplified oncogenes in a tumor-type specific fashion. To compute the significance, we assume under the null model that each amplicon interval is randomly positioned. For a genome of length *G*, the probability that amplicon *a* of length *l_a_* intersects with oncogene *g* of length *l_g_* is *A_(ag)_* = *(l_a_ + l_g_) / G.* Under the approximate assumption that all amplified intervals are independent of each other, the probability that at least 1 amplified interval in sample s randomly intersect *g* is *B*_(*s,g*)_ = 1 — ∏_*a∈s*_(1 — *A*(_*s,g*_)). As each sample is independent, the number of samples with an amplicon intersecting a particular oncogene follows a Poisson Binomial distribution with *n* trials where *n* is the number of samples with success probabilities *B*_(*1: g*)_, …, *B*_(*ng*)_. Thus, if for each tumor type with *n* samples, *g* is amplified in *k* samples, then we calculated the p-value for this observation under the null model using the PoiBin function provided by https://github.com/tsakim/poibin. To answer whether amplifications of *g* are significantly enriched in the given tumor type, we recorded an enrichment if the Bonferroni-corrected p-value was <0.05, using correction factor *G*/*l_g_* (Supplementary Figure 6).

### 13) Similarity of amplicons overlapping an oncogene

To investigate whether amplicons containing an oncogene also contained other genomic elements played a significant role in the formation or amplification process, we measured the similarity between amplicons from different samples containing one of the oncogenes EGFR, MYC and ERBB2 from the WGS dataset of 117 samples. For a given oncogene, if additional genomic elements played a significant role, then we would expect a larger overlap within amplicons from multiple samples containing the oncogene than by random chance. We measured the pairwise similarity between amplicons from 2 samples containing an oncogene, we calculated the size of the overlap between the amplicons and quantified the significance of the size of the overlap with respect to a null distribution for the size of the overlap. The null distribution was set as the distribution of size of overlap for all valid configurations for both amplicons, where a valid configuration of an amplicon was defined as any assignment of a chromosomal location for the start position of the first interval such that all the intervals maintained their order and sizes as the observed amplicon, as well as the distances between consecutive intervals such that at least 1 interval contains the given oncogene. Given a pair of amplicons, we estimated the null distribution by computing the overlap for all valid configurations obtained by shifting the amplicon intervals from the smallest to the largest possible genomic location in steps of 10000bp. Finally, for all amplicons containing each of the 3 oncogenes, we measured the significance of all pairwise similarities and visualized these through 3 QQ plots (Supplementary Fig 8). Through the QQ plots, we observed that in our sample set, that there was significant similarity within amplicons from multiple samples for any of the oncogenes. This suggests that there was no single element that played a significant role in the formation or amplification process.

### 14) Description of cervical cancer samples

We downloaded the sequencing data for 68 cervical cancer samples with matched normal samples from TCGA^23^. We downloaded 337 HPV reference genomes from PapillomaVirus Episteme database (PaVE)^32^ on Aug. 15, 2016 and concatenated these with the hg19 reference chromosomes to create an hg19_hpv337 reference genome. For each of the samples, we randomly extracted 30 million read pairs using the HTSlib bamshuf + bam2fastq utilities and aligned these to the hg19_hpv337 reference genome using BWA mem. We determined the existence and strain of HPV infecting each sample by identifying the strain with the highest number of mapped reads. For each of the identified strains, we separately concatenated the genome of the single strain to the hg19 chromosomes, to create the sample specific reference genome, for example hg19_hpv16 and mapped all the reads in the sample to this reference genome using BWA-mem. Finally, each sample, we ran AA with the -sensitivems (Methods 2) option on the mapped reads with the reference genome of the specific HPV strain as the seed interval. From the reconstructions, we combined chromosomal intervals within 5Mbp and if AA identified multiple amplified intervals connected to the HPV genome, they were treated as different amplicons unless connected to each other through discordant edges. We selected the “amplicons” for which the weighted average of the copy numbers of all amplified (CN>2.5) sequence edges from the human chromosomes was greater than 3.0. We plotted a scatter plot for the copy number vs size of all amplicons similar to Fig 2d (Supplementary Fig 9) and observed that virus-induced amplicons had a mean size of 155kbp and mean copy number of 7.04.

### 15) Identification of unifocal and bifocal signature

We developed criteria for calling a unifocal and bifocal signature and for each “amplicon” identified according to methods 13, we manually searched for unifocal and bifocal signatures. An amplicon defined to contain a unifocal signature if we found 2 reciprocal discordant edges connecting the virus genome to the human genome such that these edges had opposite strand on each of the genomes and the positions of the edges on the human genome were within 1Kbp of each other. An amplicon was defined to contain a “strong” bifocal signature if it contained a pair of chimeric edges with opposite orientations which flanked the entire amplified region on the human genome. Otherwise, an amplicon was defined to contain a “weak” bifocal signature if it contained a pair of chimeric edges with opposite orientations such that all the segments within the region flanked by these edges had higher copy number than all the other segments.

### 16) Simulation of unifocal and bifocal integrations

We hypothesized that the formation of amplicons with a unifocal signature was initiated by a unifocal integration of the virus into the human genome, whereas a bifocal integration is a more likely mechanism for the formation of amplicons with a bifocal signature. To test this hypothesis, we simulated 4 sets of amplicons where each set consisted for 40 simulations. Out of the 4 sets, 2 consisted of amplicons originating from unifocal integrations and the other 2 sets consisted of amplicon originating from bifocal integrations. For the two types of integration, we simulated one set each of linear chromosomal amplicons and circular extrachromosomal amplicons. Thus, if 4 ordered segments ABCD represent a section of the normal human genome and V represents a viral segment, then the circular and linear amplicons with unifocal integration had the initial structures [BVC] and BVC respectively. The circular and linear amplicons with bifocal integration had the initial structures [BV] and BVB respectively. Here “[ ]” represents a circular structure. For each set, the length of an amplicon, B+C in case of amplicons with unifocal integration and B in case of the amplicons with bifocal integrations, was chosen from an exponential distribution with mean 155kbp matching the mean of the amplicons detected in the 67 cervical cancer samples. In case of the unifocal distribution, the location of integration defining the segments B and C was chosen uniformly through the amplicon interval. For each simulation in all the 4 sets, we iteratively performed 20 rearrangements which was comparable to the maximum number of rearrangements in our sample set. The type of each rearrangement was chosen randomly from the set: {translocated duplication, tandem duplication, inverted duplication, translocation, inversion, deletion} with probabilities: {0.19, 0.19, 0. 19, 0.19, 0.19, 0.05} respectively and the coordinates for each rearrangement were chosen uniformly randomly from the amplicon structure formed after the previous iteration of rearrangement with the constraint that the rearrangement may not delete all segments from the viral genome. The probability of deletion was lower than the other rearrangements to make sure that too many deletions did not remove large chunks from the amplicon. After each iteration, we tested whether the amplicon structure could show a unifocal based on the existence of a pair of proximal viral connections to opposite strand of the human genome within 1kbp of each other. Similarly, we tested if the amplicon structure could show bifocal signature by checking if the virus had connection flanking the outmost ends of the human segments in the amplicon. For each set of 40 simulations and after each iteration of rearrangements, we reported the number of amplicons that showed a unifocal signature and the number of amplicons that showed a bifocal signature (Supplementary Fig 10). We found high fidelity between the type of integration and the observed signature in the amplicon. To elaborate, we found that most amplicons with unifocal and bifocal integrations showed unifocal and bifocal signatures respectively, but it was rare to observe amplicons with a unifocal integration to show a bifocal signature or amplicons with a bifocal integration to show a unifocal signature. This suggests that amplicons with bifocal signatures were unlikely to have originated from a unifocal integration of the virus, whereas amplicons with a unifocal signature were highly likely to have originated from a unifocal integration.

